# Seasonal dynamics of nitrification in the water column of marine Lake Grevelingen

**DOI:** 10.1101/2025.06.29.662106

**Authors:** Nicky Dotsios, Jessica Venetz, Olga M. Zygadlowska, Wytze K. Lenstra, Niels A.G.M. van Helmond, Anna J. Wallenius, Robin Klomp, Caroline P. Slomp, Mike S.M. Jetten, Maartje A.H.J. van Kessel, Sebastian Lücker

## Abstract

Coastal ecosystems serve as vital connectors between land and ocean, and their nitrogen cycle and ammonium removal can be affected by various factors. During our seasonal sampling campaign in 2021, the eutrophic marine Lake Grevelingen exhibited high ammonium concentrations and low nitrification rates in the water column, except for a brief period in late summer. Our study revealed ammonium accumulation in the anoxic middle and bottom water layers due to restricted transport caused by water column stratification. Only when ammonium reached the oxic part of the water column, was a short-lived peak in nitrification activity and nitrifier abundance observed at the end of August. Amplicon sequencing indicated very low abundances of *Nitrosococcus* and *Nitrospira* (<0.2%) in March, while *Nitrosomonas* and *Nitrospina* peaked at the end of August with relative abundances of 2.5% and 1.3%, respectively. Archaea and archaeal ammonium oxidizers were found in very low abundances. Anammox 16S rRNA genes were not detected. Together, these observations suggest a limited role for nitrification in ammonium removal in marine Lake Grevelingen.

## Introduction

Coastal marine ecosystems are highly dynamic environments significantly influenced by human activities (Breitburg et al., 2018). These systems are often characterized by substantial nutrient loading and organic matter deposition, which can result in oxygen depletion and the formation of hypoxic or anoxic zones (Breitburg et al., 2018; Diaz & Rosenberg, 2008). The severity and duration of coastal hypoxia are impacted by seasonal water column stratification, which arises from temperature- and salinity-driven density gradients that inhibit vertical mixing (van Haren, 2019). Stratification typically intensifies during summer as temperatures rise, limiting oxygen replenishment to bottom waters, while mixing in fall or winter restores oxygen levels. Oxygen availability is crucial in regulating nitrogen cycling processes in coastal marine environments. When high concentrations of ammonium enter the water column, its removal by nitrification requires sufficient oxygen. Therefore, understanding how oxygen availability influences nitrification is essential when assessing ammonium removal and nitrogen cycling in coastal environments.

Nitrification, the microbial oxidation of ammonia (NH_3_) to nitrate (NO_3_^-^) via nitrite (NO_2_^-^), is an important process in the marine nitrogen cycle (Hutchins & Capone, 2022). It is carried out by various groups of microorganisms. The first step, ammonia oxidation, is performed by ammonia-oxidizing bacteria (AOB) and archaea (AOA), which oxidize NH_3_ into NO_2_^-^ (Hutchins & Capone, 2022). Subsequently, nitrite-oxidizing bacteria (NOB) convert NO_2_^-^ into NO_3_^-^ (Daims et al., 2016). Comammox *Nitrospira* bacteria can perform the complete oxidation of ammonium to nitrate (Daims et al., 2015; Van Kessel et al., 2015). Traditionally, nitrification was considered a strictly aerobic process; however, recent studies have shown that it can also occur in oxygen-deficient zones (Bristow et al., 2016; Molina et al., 2010) and coastal marine water columns (Bristow et al., 2015; Jäntti et al., 2018), suggesting that certain nitrifying microorganisms are adapted to low oxygen conditions or possess alternative metabolic pathways.

In marine systems, AOA - particularly those of the genus *Nitrosopumilus* - are often more abundant than AOB as they are typically more competitive in low ammonium environments (Bouskill et al., 2012; Francis et al., 2005; Lam et al., 2007; Newell et al., 2013; Santoro et al., 2010). AOB, while less prevalent in marine systems, have been identified in some estuaries and include genera such as *Nitrosospira* (Bouskill et al., 2012; Freitag et al., 2006), *Nitrosococcus* (Ward & O’Mullan, 2002), and *Nitrosomonas* (Bollmann & Laanbroek, 2002; De Bie et al., 2001). NOB, primarily those affiliated with the phylum *Nitrospinae*, dominate marine environments (Bristow et al., 2015; Damashek & Francis, 2018; Füssel et al., 2012; Mincer et al., 2007; Sun et al., 2019), while certain marine *Nitrospira* species can also play a significant role (Nunoura et al., 2015).

In addition to nitrification, anaerobic ammonium oxidation (anammox) plays a important role in nitrogen removal in oxygen-deficient zones (Jensen et al., 2008; Kalvelage et al., 2011; Kuypers et al., 2003, 2005). In marine systems, *Scalindua* is the dominant anammox bacterium (Van de Vossenberg et al., 2013); however, two metagenomes of new marine anammox species, *Candidatus* Anammoxibacter and *Candidatus* Bathyanammoxibiaceae, have been recently described (Suarez et al., 2022; Zhao et al., 2024).

While many studies have focused on nitrogen dynamics and ammonium removal in (near) permanent oxygen-deficient zones in the ocean (Frey et al., 2014; Kuypers et al., 2018; Lam & Kuypers, 2011) , less is known about the controls on ammonium removal in seasonally stratified oxygen-deficient coastal marine systems.

In this study, we therefore investigated the dynamics of nitrification in the water column of marine Lake Grevelingen, a eutrophic coastal system in the Netherlands, which experiences seasonal stratification in its deeper basins, resulting in hypoxic and anoxic bottom waters during the summer (Venetz et al., 2023; Żygadłowska et al., 2023). The microbial ecology of marine Lake Grevelingen is complex and influenced by various factors, including oxygen and nutrient availability (Venetz et al., 2023). While marine anammox bacteria have been studied in the sediments of the Den Osse basin of Lake Grevelingen (Lipsewers et al., 2016), their activity and contribution to nitrogen removal in the water column remain poorly understood. During a seasonal sampling campaign in 2021, we focused particularly on the distribution and activity of nitrifying microorganisms at different water depths. By exploring the interplay between nitrification, stratification, and oxygen availability, we aimed to enhance our understanding of ammonium removal and nitrogen cycling in this seasonally stratified eutrophic marine system and its implications for nitrogen removal.

## Material and Methods

### Fieldwork location

Lake Grevelingen, located in the southwest of the Netherlands (Fig. S1), was formed in the 1970s by the construction of two dams. By regularly opening the sluices towards the North Sea, a marine salinity ranging from 29 to 33 psu is maintained. The Scharendijke basin (51.742°N, 3.849°E) reaches a depth of 45 meters and is the deepest part of the lake. It experiences seasonal stratification, which leads to hypoxic and anoxic bottom waters in late summer. More detailed descriptions of the study site can be found elsewhere (Egger et al., 2016; Żygadłowska et al., 2023) During multiple sampling campaigns from March to October 2021, and in July 2022, September 2023, and August 2024 with the R/V Navicula (see Table S1 for specific dates), the biogeochemistry (as described in Żygadłowska et al., 2024), microbial diversity, and nitrification rates in the water column of the Scharendijke basin were studied.

### Water column sampling

Conductivity, temperature, and depth were determined with a CTD unit (SBE 911 plus, Sea-Bird Scientific, Bellevue, WA, USA). The oxygen concentrations were measured with a SBE43 Dissolved Oxygen Sensor (Sea-Bird Scientific). Water samples were collected with 10 L Niskin bottles at 1 or 2 m intervals up to 43 m depth. Unfiltered water was collected for DNA extraction in 1 L pre-rinsed plastic bottles at 30 discrete depths. For nitrification rate measurements, unfiltered water was collected in 1 L Schott bottles. The bottles were rinsed with water from the depth of interest, completely filled, and stored in the dark at 4°C until further processing. In the following years (July 2022, September 2023, and August 2024), an adapted nitrification activity measurement method was utilized to minimize oxygen contamination. 120 ml serum bottles were filled directly from the Niskin bottle using Norprene tubing, ensuring that the water overflowed the serum bottles at least three times to minimize air entrapment. The bottles were sealed with red butyl stoppers and metal caps, checked to confirm the absence of air bubbles, and stored in the dark. Water samples were collected and sulfide, ammonium, nitrite, and nitrate levels were measured using colorimetric assays as detailed by Żygadłowska et al. (2024).

### Upward mixing and accumulation of NH4+ in the water column

To determine whether upward mixing of NH_4_^+^ released from the sediment could explain the vertical distribution of NH_4_^+^ in the water column between June and September 2021 and in July 2022, September 2023 and August 2024, we calculated NH_4_^+^ concentrations (*C*) as a function of water depth (*z*) assuming upward vertical transport through turbulent diffusion only (Lewis & Landing, 1991):

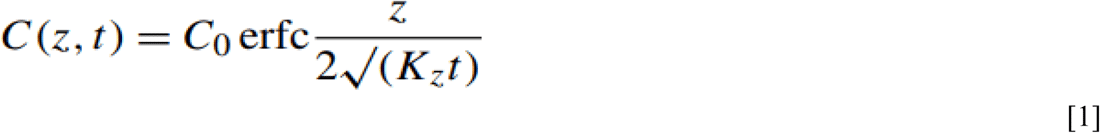

where *C_o_* stands for the bottom water NH_4_^+^ concentration, *t* is the duration of stratification at the time of sampling, and *K_z_* is the vertical eddy diffusivity. The *K_z_* profile was estimated from the density profile, inferred from the temperature and salinity measurements, following the procedure described in Żygadłowska et al. (2023).

Water column concentrations of NH_4_^+^ were integrated with depth to estimate the total amount of NH_4_^+^ in the water column (in mol m^-2^) for each sampling event in 2021. Three vertical sections of the water column were considered: the upper (0–10 m), middle (10–35 m), and bottom (35–45 m) layers.

### Potential ammonium and nitrite oxidation rates

Water samples for nitrification rates were collected at five discrete depths throughout the water column during mid-August, the end of August, September, and October in 2021 (Table S1). Depths were chosen based on the CTD oxygen and temperature profiles, targeting the oxic zone, the start and middle of the oxycline, the oxic-anoxic interface, and the anoxic zone. Water samples were collected as described above, transported to the lab, and nitrification rate measurements were started the following day. Water (100 ml) was transferred into sterile, autoclaved 120 ml borosilicate serum bottles. For oxic conditions, the bottles were sealed with a gray bromo-butyl stopper, while for anoxic conditions, they were sealed with red butyl stoppers and capped. For both aerobic and anaerobic incubations, separate incubations with 500 µM NH4+, 500 µM NO2-, or no substrate addition were conducted. In the adapted “overflow” method, the serum bottles were amended with the respective substrate immediately after sampling. First, 20 ml of water was removed with a syringe by adding an overpressure of an Argon/0.5% CO2 mixture with a second syringe. The final volume was 100 ml of water with approximately 1 bar overpressure and 100 µM NH4+, 50 µM NO2-, 100 µM NH4+ + 50 µM NO2-, were added. Subsequently, 1 ml samples were taken with a gastight syringe at 0, 2, 4, 6, 24, 48, 72, and 168 hours. Liquid samples were filtered through 0.2 µm nylon syringe filters (Arcodisc®, Pall Corporation, City, Country) and stored at −20 °C until further analysis.

### Colorimetric assays

Ammonium was measured using a modified colorimetric *o*-phthaldehyde assay (Roth, 1971) by mixing 10 µl of sample with 150 µl reagent containing 4 mM *o*-phthaldehyde, 1% (v/v) ethanol and 0.005 β-mercaptoethanol in 400 mM K_x_H_x_PO_4_ (pH 7.3), 10% (v/v). After incubation for 20 min incubation in the dark, the absorbance was measured at 405 nm using a Tecan Spark M10 plate reader (Tecan Group Ltd., Männedorf, Switzerland). Nitrite and nitrate were quantified using a modified Griess assay by mixing a 1:1 mixture of reagents A and B with 1 volume of sample. Reagent A consists of 1% sulfanilic acid in 1 M HCL, and reagent B is a 0.1% (w/v) N- (1-naphtyl) ethylenediamine solution. After incubation for 10 minutes and the absorbance was measured at 540 nm on a SpectraMax190 plate reader (Molecular Devices, San Jose, US). To measure nitrate in the same sample, nitrate was reduced to nitrite by adding 6.9 mM VCl_3_ and incubation for 30 min at 60 °C. Absorbance was then remeasured at 540 nm (Griess, 1879; Wang et al., 2016).

### Analysis of 16S rRNA gene amplicon sequencing data

In this study, we focused on analyzing the diversity of nitrifying microorganisms using the water column 16S rRNA gene amplicon sequencing data available at the National Center for Biotechnology Information (NCBI) under accession number PRJNA1053269, published by Venetz et al. (2024) and the sediment 16S rRNA gene amplicon sequencing data available under accession number PRJNA1257658. Quality control and trimming of the sequencing data were performed using the DADA2 pipeline in RStudio (Callahan et al., 2017). The phyloseq package (McMurdie & Holmes, 2013) was used for relative abundance calculations. Data visualization was performed with *ggplot2* (Wickham, 2016). To obtain comparable estimates of bacterial and archaeal relative abundances in the water column and the sediment of the Scharendijke Basin, we used SingleM (Woodcroft et al., 2024) on raw metagenomic reads obtained from Venetz et al. (2023) and Wallenius et al. (2024) deposited at the European Nucleotide Archive under accession numbers PRJEB57287 and PRJNA1167897 (Biosample numbers SAMN47278200 -SAMN47278204), respectively.

### Differential abundance and network analysis

Differentially abundant bacterial families between sampling months were identified using the run_aldex() function from the ALDEx2 R package (Fernandes et al., 2014). Count data were normalized with the rarefaction method to account for varying sequencing depths across samples. Statistical analysis was performed using the Wilcoxon rank-sum test to evaluate differences in microbiome composition in the full water column during the following two time periods: July *vs*. August and August *vs*. end of August. The Benjamini-Hochberg method was applied to adjust p-values for multiple comparisons, with a false discovery rate threshold of < 0.05. The effect size was then illustrated as a bar plot using ggplot2. To investigate the correlation between microbial community members and clarify differences between the oxic (> 60 µM O_2_), hypoxic (< 60 µM O_2_), and anoxic (< 4 µM O_2_ and presence of sulfide) water layers in July, August, and end August, we conducted a network co-occurrence analysis using the NetCoMi package (Peschel et al., 2021). To capture the diversity of dominant taxa in each water layer, Pearson correlations were calculated for the 30 most abundant taxa. The “signed” distance metric was employed to convert correlations into dissimilarities in a manner that emphasizes strong negative relationships as greater distances. For sparsification, a correlation threshold of > 0.3 was applied before transformation into the adjacency matrix. To address the compositional nature of the sequencing data, read counts were transformed into Euclidean space via centered log-ratio (clr) transformation to mitigate compositional effects. Clusters were identified using the cluster_optimal algorithm, and centralities were calculated through eigenvector decomposition. Hubs were defined as betweenness centrality, where nodes are characterized by centrality values above the 95^th^ percentile. Cluster connectivity was quantified as the ratio of between-cluster edges to within-cluster edges.

## Results

### Water column chemistry

The seasonal sampling in 2021 at the Scharendijke basin in Marine Lake Grevelingen revealed both temporal and spatial differences in the nutrient depth profiles. From March to May, the water column was fully mixed, oxic, and contained nitrate (Fig. 1, Fig. S2). In March, NH_4_^+^ concentrations in the bottom water were relatively low, reaching a maximum of 9 µM, but began to increase in April and May, reaching 29 µM and 54 µM, respectively. From June onwards, stratification was observed, accompanied by increased H_2_S concentrations (54 µM) in the bottom water (Fig. 1). At the same time, increased amounts of NH_4_^+^ began to diffuse from the sediment into the overlying water, with concentrations rising to 395 µM in the bottom water.

**Figure 1.**
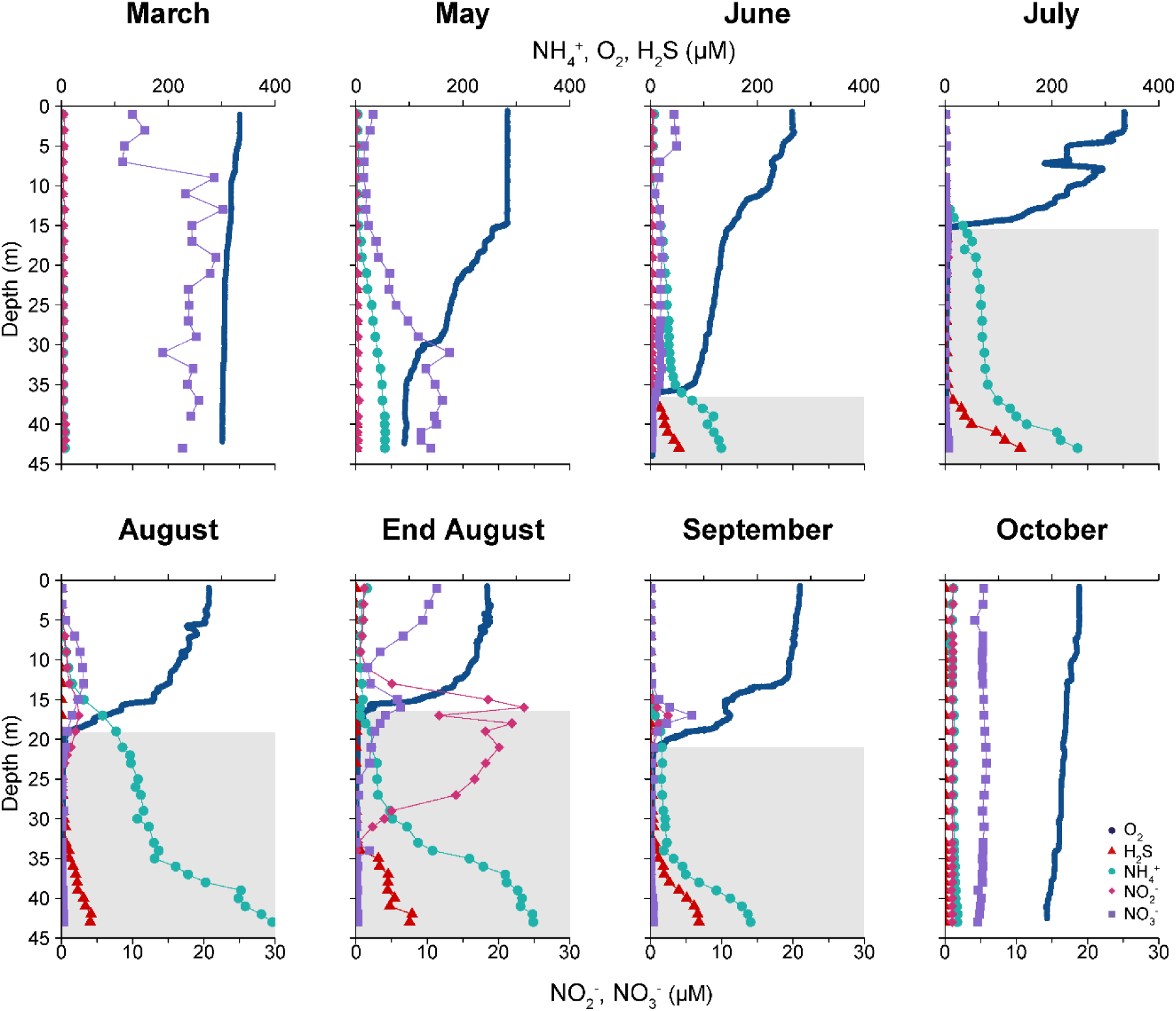
Water column profiles of the Scharendijke basin in Lake Grevelingen in 2021. The depth profiles of O_2_ (blue circles), H_2_S (red triangles), NH_4_^+^ (green circles), NO_2_^-^ (pink diamonds), and NO_3_^-^ (purple squares) are depicted. The grey shaded areas indicate an O_2_ concentration that was below the detection limit of the sensor (< 4 µM).

From July onwards, a suboxic zone (anoxic, without H_2_S) developed, which persisted until September. Below 30 m, H_2_S concentrations increased over time, reaching a maximum of 140 µM in July. In September, the H_2_S concentration decreased to 91 µM; no H_2_S was present in the water anymore in October. The suboxic zone was not observed in the following years (Fig. S2) (Żygadłowska et al., 2024). Between July and August, the oxycline shifted downward, allowing

NH_4_^+^ to reach the oxic water column, with NH_4_^+^ concentrations of 34 µM in July and 102 µM in August. By the end of August, NH_4_^+^ concentrations had decreased in both the oxic and suboxic zones, coinciding with a rise in nitrite levels (19.7 µM) around the oxycline, suggesting microbial ammonia oxidation (Fig. 1). This NO_2_^-^ peak was still observed in September but had decreased substantially (2.1 µM; Fig. 1). In the following years, we tried to sample at the end of the summer period to assess whether similar phenomenona recurred. While we observed NH_4_^+^ reaching the oxic part of the water column in July 2022, we did not detect nitrification in the summers of 2022, 2023, and 2024 (Fig. S2).

### Vertical transport and accumulation of NH_4_^+^

Modelled NH_4_^+^ concentrations, which assume upward turbulent mixing of NH_4_^+^ from the bottom water only, are generally lower than the measured water column NH_4_^+^ concentrations (Fig. 2; Fig. S3), suggesting an additional source of NH_4_^+^. Integrated water column concentrations of NH_4_^+^ for 2021 indicate an accumulation of NH_4_^+^, primarily in the **middle layer** (10–35 m) and bottom water layer (35–45 m) of the water column, following the onset of stratification (Table S2). By the end of August, NH_4_^+^ concentrations in the middle layer decreased, suggesting that an active removal process was occurring. This decrease coincided with the appearance of NO_2_^-^ and NO_3_^-^ (Fig. 1), suggesting activity of nitrifiers. From this time onwards, the water column stratification began to weaken until the water column became fully mixed again in October (Fig. 1; Fig. S4; Żygadłowska et al., 2024). Concentrations of NH_4_^+^ remained lower in the middle and bottom water layers at the sampling times in the subsequent years (July 2022, September 2023, and August 2024; Fig. S3), which may be the result of differences in water column stratification and/or NH_4_^+^ input from the sediment; Fig. S4).

**Figure 2.**
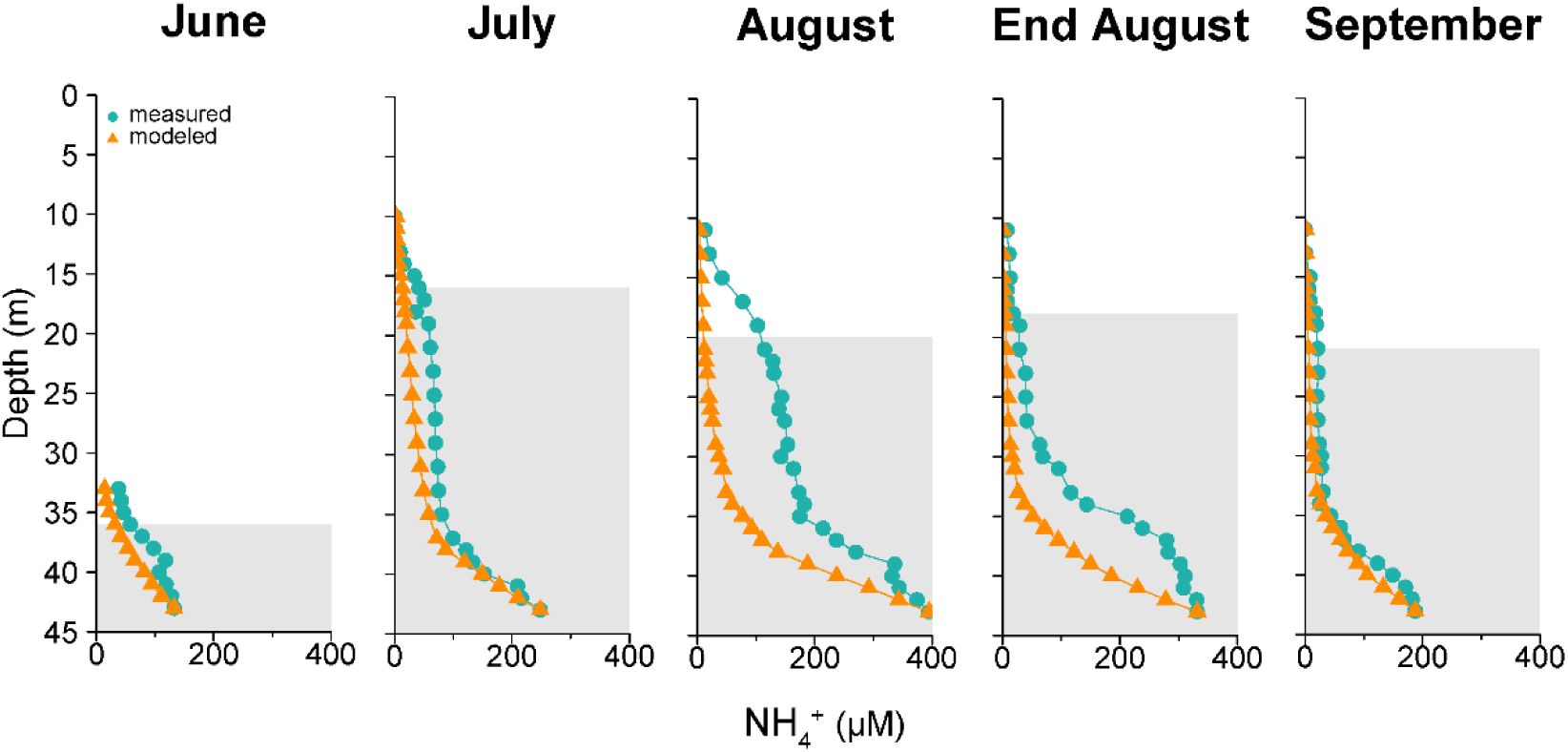
Comparison of the measured NH_4_^+^ depth profiles in the water column during the sampling campaigns of 2021 and the predicted NH_4_^+^ concentrations based on the turbulent diffusion model, calculated for the stratified part of the water column. The measured NH_4_^+^ concentrations are indicated by green circles, while orange triangles indicate the modelled NH_4_^+^ concentrations.

### Water column nitrification rates

We tested for nitrification at five distinct depths throughout the water column from August to October 2021. Consistent with the chemical nutrient profiles, the highest ammonia oxidation rates were observed near the oxic-anoxic interface, reaching 51 µM/day at 21 m depth at the end of August, and 34 µM/day at 13 m depth in September (Fig. 3).

**Figure 3.**
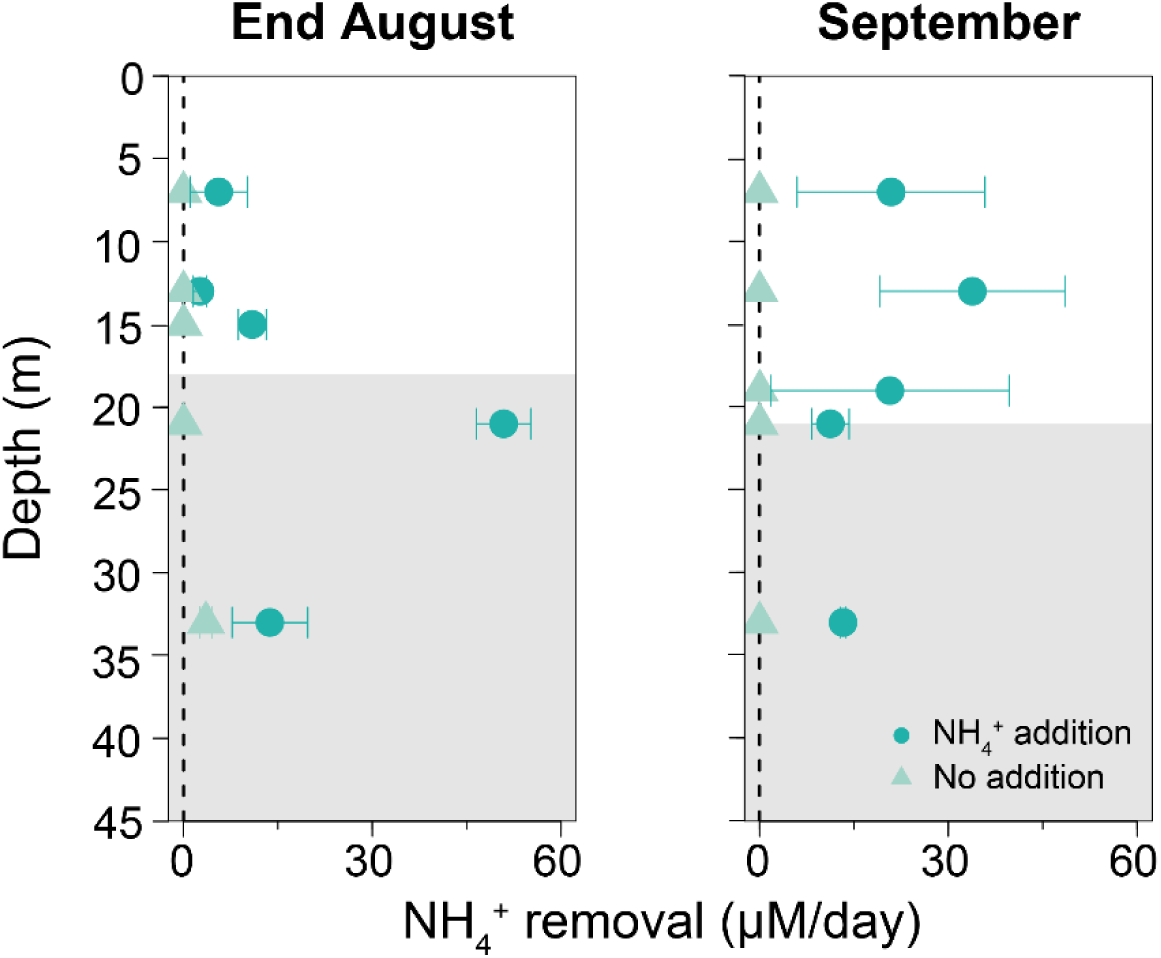
Depth profiles of NH_4_ ^+^ removal rates at selected time points at the End of August and September. The NH_4_ ^+^ removal rates (µM/day) are shown for the incubations with (dark blue) and without NH ^+^ (only endogenous NH_4_ ^+^, light blue). Data points represent the mean ammonium removal rates of duplicate measurements for 5 selected depths. The error bars represent ± standard deviation (SD) based on duplicate measurements. A dashed black line at 0 µM/day indicates the threshold of net ammonium removal (positive values). The shaded areas indicate the water depths where O_2_ was below the detection limit of the sensor (< 4 µM).

In the surface water at 5 m depth, ammonia oxidation rates of 7 µM/day were observed in August. In October, no activity was observed in the oxic water layers, and even a small NH_4_^+^ increase was observed (Fig. S10). Under strictly anoxic conditions, no activity was observed. Interannual variability was evident, with less than 4 µM/day NH_4_^+^ removal recorded at all depths in July 2022, September 2023 and August 2024.

### 16S rRNA gene sequencing-based seasonal variability of the microbial community

We investigated the relative abundance of nitrifying bacteria using 16S rRNA gene amplicon sequencing. Nitrifiers of the families *Nitrospiraceae* and *Nitrosococcaceae* were detected at very low relative abundances (< 0.2 %) in March but not in any other month in 2021 (Fig. S5). *Nitrosomonadaceae* and *Nitrospinaceae* abundances were low in March (< 0.15 and 0.03 %) but increased to 2.5 % and 1.3 %, respectively, by the end of August (Fig. 4).

**Figure 4.**
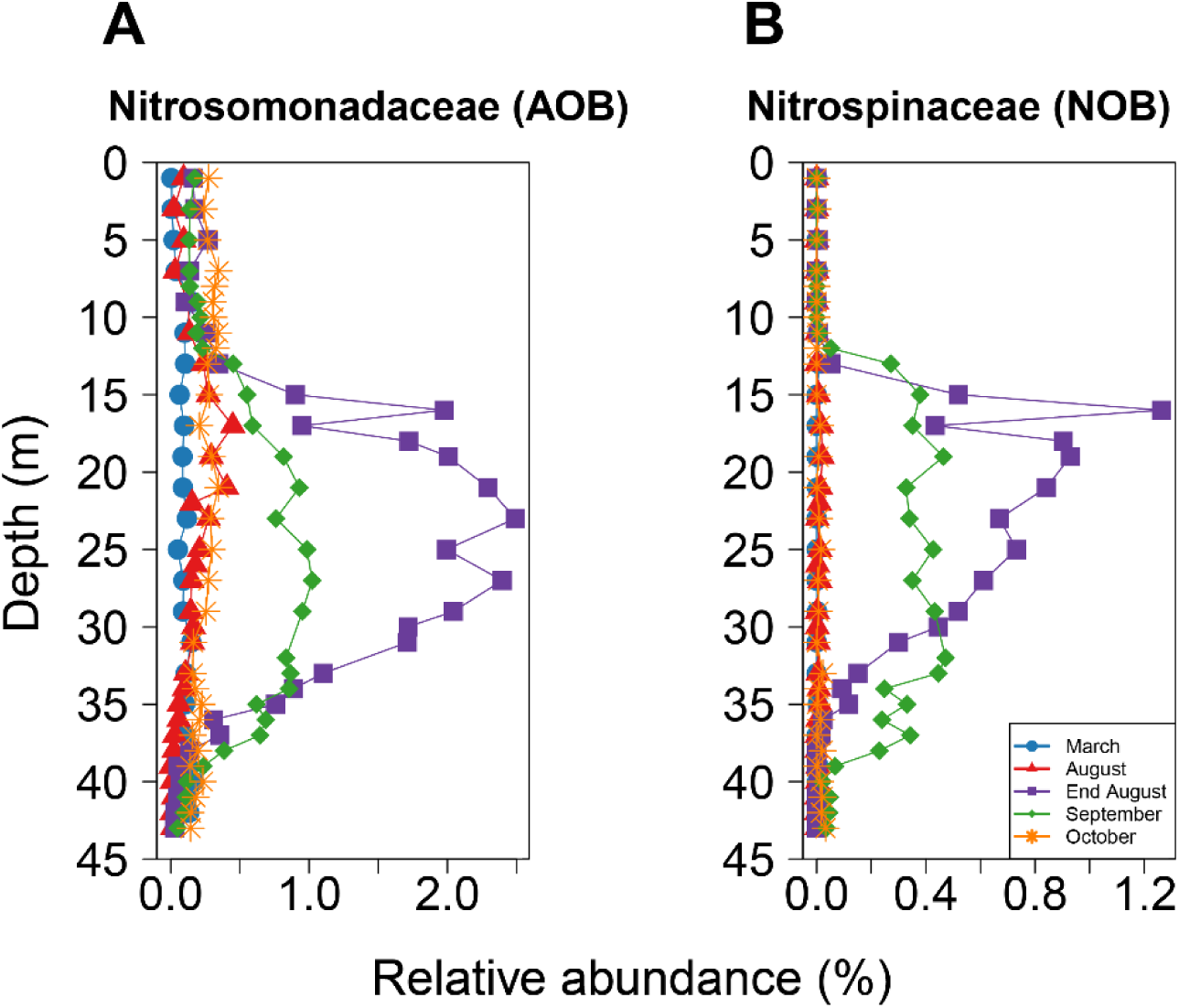
16S rRNA gene sequencing-based nitrifier abundance in the Scharendijke water column of Lake Grevelingen in 2021. Relative abundances of (A) the ammonia oxidizing Nitrosomonadaceae and (B) the nitrite oxidizing Nitrospinaceae families in March (blue circles), August (red triangles), End August (purple squares), September (green diamonds), and October (orange stars).

In September, the relative abundances of these nitrifiers started to decrease again and were below 0.3 % by October. Based on metagenomic read classification, the total archaeal fraction in the water column of the Scharendijke basin was 3.3 % in September 2020 (Table S3). Ammonia-oxidizing archaea of the family *Nitrosopumilaceae* were found in the water column, especially at the oxic-anoxic interface, and made up to 6.3 % of the archaeal fraction in July 2021, but were also present below this zone (Fig. S7). In other months, the relative abundances of archaea were between 0.4 and 2.6 % (Fig. S7). In contrast, the *Nitrosospumilaceae* had a relative abundance up to 48% of the archaeal fraction (and Archaea were 2% of all reads in SingleM analysis; Table S4) in the sediment (Wallenius et al., 2024), indicating they could be involved in nitrate production during the oxic winter months (Fig. S8). Intriguingly, in the summer period, the relative abundance of *Nitrosopumilaceae* in the top sediments declined to below 6 % (Wallenius et al 2024). A similar seasonal pattern was observed for bacterial nitrifiers in the sediment. They were also more abundant in top sediments during winter compared to summer (Fig. S9.

### Differential abundance and co-occurrence analysis

To track the changes in microbial families from month to month, we performed a differential abundance analysis of the complete bacterial community throughout the summer. Notable increases from July to August were seen for the *Flavobacteria* and *Pseudomonas* families (positive effect size), while the most significant decline was recorded in SAR324 and the *Marinimicrobia* (negative effect size; Fig. 5A). From early August to the end of the month, *Rhodothermaceae*, *Arinelleaceae*, SAR406, and *Nitrosomonadaceae* showed the largest increases, whereas *Proxilibacteriaceae, Cyclibacteriaceae,* and PB19 experienced the most considerable decline (Fig. 5B).

**Figure 5.**
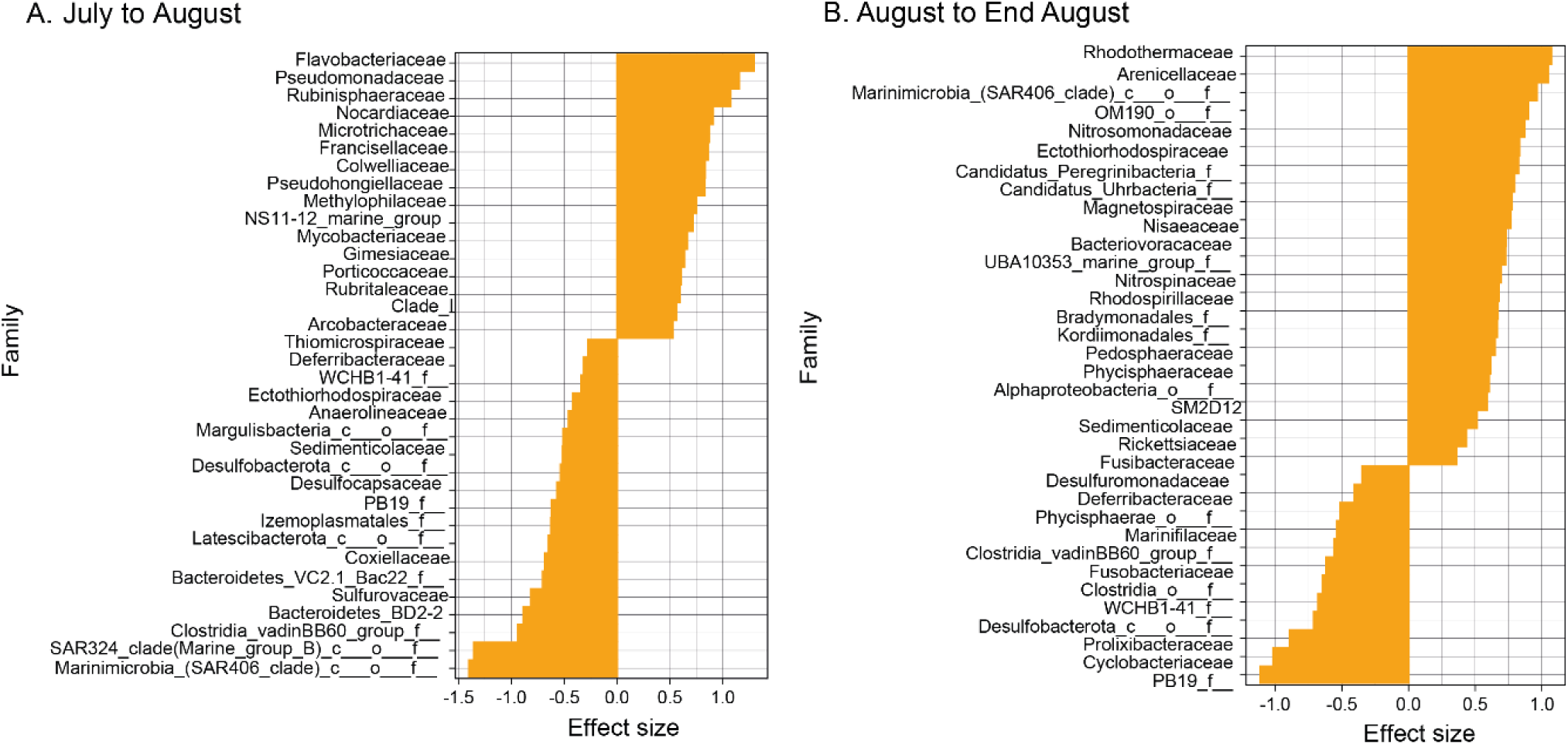
Effect sizes of differentially abundant bacterial families between July and August (A) and August and end August (B). The effect size was computed from the 30 most abundant families based on the Wilcox Test (HB corrected p-value < 0.05, n = 30). Positive values indicate increased abundance at the second time point included in the analysis, while negative values indicate higher abundances at the first.

We examined the potential interactions of *Nitrosomonadaceae* with other community members through co-occurrence network analyses. As *Nitrosomonadaceae* showed a strong increase in abundance in end August, and had a low abundance in the other months, we focused our analysis on the microbial community in July and August. Network analysis revealed strong connections with the families OM190, SAR202, and *Candidatus* Peregrinibacteria (Fig. 6). When comparing the co-occurrence networks of July and August, we observed that the *Sulfurimonas* species were important in the hypoxic zone in July and August but disappeared by the end of August (Fig. 6; Fig. S12).

**Figure 6.**
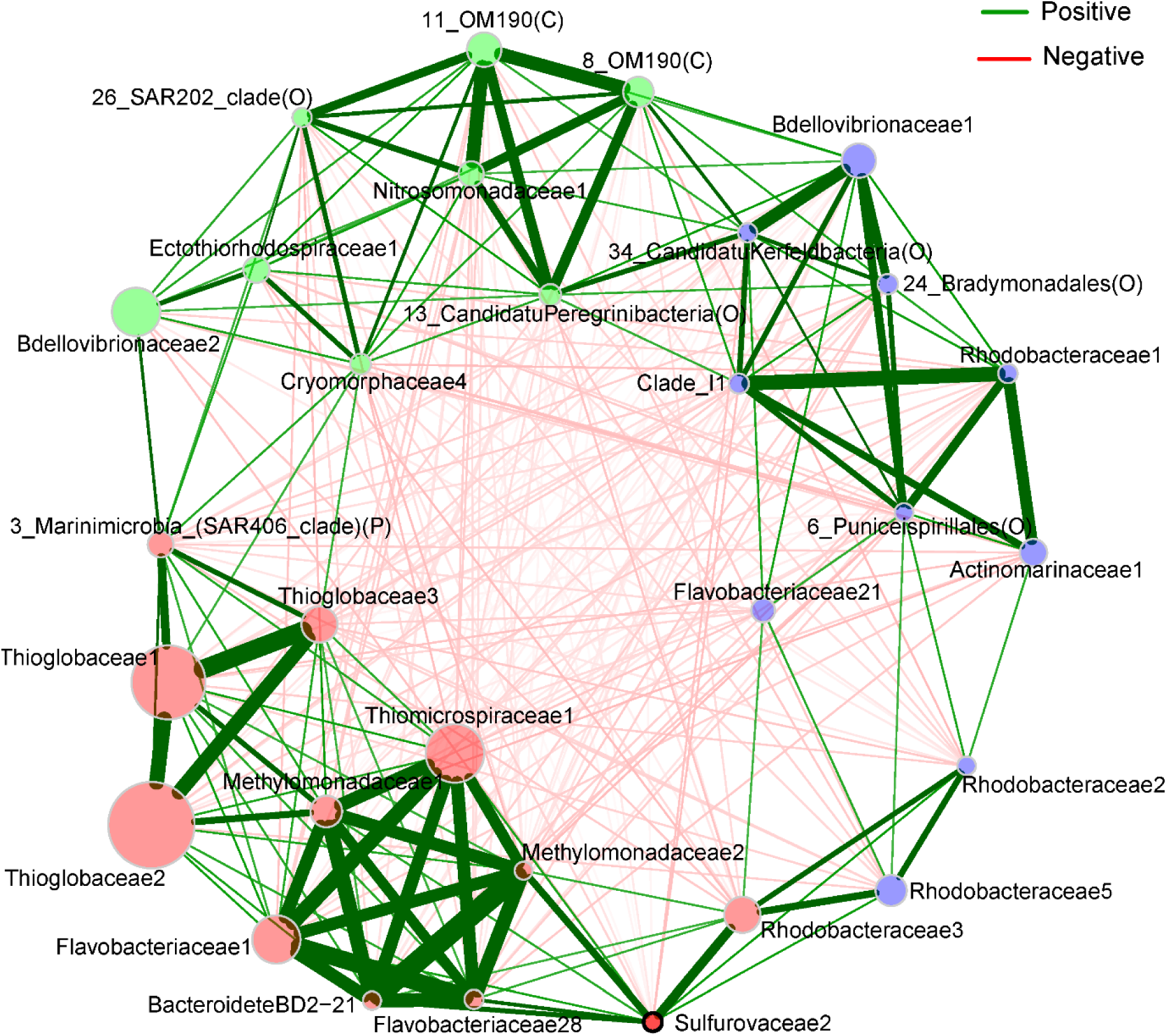
Co-occurrence networks based on Pearson correlation for the hypoxic (60 µM O_2_) zone in the water column at the end of August. Edges represent significant correlations, with green lines indicating positive correlations and red lines negative correlations. Line width represents the strength of the positive correlations. Node colors indicate distinct microbial clusters, while node size represents relative abundance. Hubs, identified based on betweenness centrality, are highlighted with black node borders and bold font and represent key microbial taxa that serve as connectors between different clusters.

## Discussion

This study examined the NH_4_^+^ turnover dynamics in Marine Lake Grevelingen. The deepest basin, Scharendijke, serves as a representative site for investigation due to its seasonal stratification and elevated NH_4_ ^+^ levels. Nonetheless, our research - utilizing high-resolution chemical profiles, potential nitrification assays, and 16S rRNA abundances - reveals that nitrification is only significant in this system when NH_4_ ^+^ reaches the oxic-anoxic interface of the water column.

### NH_₄_^+^ dynamics in the water column

Throughout the summer months, NH_4_ ^+^ was consistently present in the bottom and middle layers of the stratified water column of Scharendijke basin (Fig. 1; Fig. S2; Fig. S4). Water column stratification, a well-known feature in this lake (van Haren, 2019), strongly limited vertical water and solute transport, including that of NH_4_ ^+^. The difference between the measured and modelled NH_4_ ^+^ concentrations in the water column (Fig. 2) points to a possible additional source of NH_4_ ^+^. Given the bathymetry of the lake, with several deep basins located within an overall shallower system (Fig. S1), this source could be the lateral input of NH_4_^+^ from surrounding shallower areas (Fig. S1). Notably, modeled and measured NH_4_ ^+^ profiles were quite similar in 2022 and 2023 (Fig. S3), suggesting a more stagnant water column at the time of sampling.

From July to August, the oxycline deepened, allowing the zones where NH_4_ ^+^ and oxygen were present to overlap. From the end of August onwards, a decrease in NH_4_ ^+^ concentrations in the middle layer (10-35 meter) coincided with an increase in concentrations of NO_2_^-^ and NO_3_^-^ in the same layer, suggesting NH_4_ ^+^ removal through microbial nitrification. From this time onwards, reduced density gradients and changes in profiles of a range of solutes point towards a weakening of stratification (Żygadłowska et al., 2024), which likely contributed to enhanced vertical transport of NH_4_ ^+^ within the system. By October, the water column was fully mixed, as is evident from the uniform density profile across the water column. Still, NH_4_ ^+^concentrations remained higher than those in March (< 7 µM; Fig. 1, S2), suggesting a further decrease occurred during the winter months. During these months, most of the ammonium is presumably oxidized to nitrate in the oxic sediment layers by *Nitrosopumilus* and NOB, which constitute a significant portion of the archaeal fraction in the sediment (Fig. S8; Fig. S9; Rigutto et al., 2025; Wallenius et al., 2024). Nitrate then accumulates in the water column which remains fully mixed in winter, explaining the even distribution of nitrate with water depth in March.

### Dynamics of nitrification

Water column chemistry indicates that microbial ammonia oxidation occurs almost exclusively at the end of August and in September. This is corroborated by the elevated potential NH_4_ ^+^ removal rates measured during these months, while minimal removal or even NH_4_ ^+^ accumulation was observed in other periods. Depth profiles reveal that NH_₄_^+^ reaches the oxic part of the water in both July and August, potentially inducing growth of nitrifiers and consequently the nitrification activity observed by the end of August and September. The downward shift of the oxycline in August further enlarged the zone with availability of both NH_4_ ^+^ and oxygen, thus promoting nitrification. Taken together, this suggests that in 2021, microbial ammonium oxidation only occurred in late August and September, consistent with the 16S rRNA gene sequencing-based results showing an increase in *Nitrosomonadaceae* and *Nitrospinaceae* abundances close to the oxic-anoxic interface at the end of August. Their approximately four-fold increase in relative abundance within two weeks (mid-August to end-August) indicates rapid growth of these nitrifiers when the conditions are favorable.

Although the nitrification rates, especially in estuaries can range from < 1 nM day^−1^ to 100 µM day^−1^; (Damashek et al., 2016; Rasmussen & Francis, 2024), the ammonium removal rates at the oxic-anoxic interface in the Scharendijke basin were notably high, reaching 51 and 34 µM/day at the end of August and in September, respectively (Fig. 3). In contrast, the maximum ammonia oxidation rates recorded in similar system like the San Francisco Bay (NH_4_^+^ < 12 µM) _and_ the Bornholm Deep (NH_4_^+^ < 4 µM) in the Baltic Sea were 1.8 and 0.88 µM/day, respectively (Berg et al., 2015; Rasmussen & Francis, 2024). One study conducted in the Gulf of Mexico (NH_4_ ^+^ < 3 µM) documented nitrification rates of 3.5 µM/day (Carini et al., 2010), which is still nearly ten times lower than the rates observed in the Scharendijke basin. However, in these systems, *Nitrosopumilus* was the primary ammonia oxidizer, which is associated with lower conversion rates. In other years, ammonia oxidation and the concomitant nitrite peak were not observed, with NH_4_ ^+^ removal rates approaching zero or even some net NH_4_ ^+^ production. Although ammonia oxidation appears relatively quickly at the end of August, it lasts for only a short duration, suggesting that the nitrifiers in Scharendijke basin can effectively remove NH_4_ ^+^ when favorable conditions, i.e., the co-occurrence of both NH_4_ ^+^ and oxygen, are present.

The observed NH_4_^+^ removal coincides with NO_2_^-^ production (Fig. 1). However, the observed NO_2_^-^ concentrations do not match the consumed NH_4_^+^ which suggests removal of the NO_2_^-^ via NOB or denitrification (Rigutto et al., 2025). This substoichiometric nitrite production was also observed in the activity assays (Fig. S11). For the NO_2_^-^ oxidation, a similar seasonal pattern is observed for the ammonium removal. At the end of August, there is a peak in NO_2_^-^ production, which implies that NO_2_^-^ oxidation responds more slowly than ammonium oxidation to favorable conditions. This has also been reported in the water column of the Gulf of Mexico, where ammonia oxidation was 30 times faster compared to NO_2_^-^ oxidation (Bristow et al., 2015). Both at the end of August and in September, NO_3_^-^ production was observed in the Scharendijke basin, indicating that nitrite oxidation is occurring, which is corroborated by the increased relative abundance of *Nitrospinaceae* in the 16S rRNA gene sequencing data.

### Minor contribution of nitrification in oxygenated surface water

The observed increase in NH_4_^+^ concentrations in the water column of Marine Lake Grevelingen, coupled with the low abundance of nitrifiers throughout most months during our sampling campaign in 2021, suggests a minor contribution of nitrification in the Scharendijke basin. During spring, when the water column is oxygenated, the relative abundance of nitrifiers remains low while ammonium concentrations increase, indicating that nitrification in the water column plays a minimal role during this period. However, in summer, ammonia oxidation becomes somewhat active in oxic surface waters, although this is considerably less pronounced compared to the oxic-anoxic interface. In other ecosystems, such as the Gulf of California, the Eastern Tropical North Pacific Ocean (Beman et al., 2012), Sargasso Sea (Newell et al., 2013), Arabian Sea (Newell et al., 2011), and Ems Estuary (Sanders & Laanbroek, 2018), nitrifiers are found throughout oxic water columns, as evidenced by bacterial or archaeal *amoA* gene qPCR copy number presence. In these systems, nitrifier populations appear to peak at or just below the euphotic zone (Santoro et al., 2010; Zakem et al., 2018). However, in the highly eutrophic freshwater Onondaga Lake, no significant nitrification rates were observed in the water column despite high concentrations of ammonium and oxygen (Pauer & Auer, 2000). This lake exhibits irregular nitrification rates over a 7-year sampling span (Gelda et al., 2000). This indicates that systems with high NH_4_^+^ concentrations do not necessarily support biological NH_4_^+^ removal. It is crucial to understand and conduct further research on why nitrifiers are absent in these systems to prevent future nitrogen build-up and eutrophication, which could lead to more deoxygenated waters.

Along with the low abundances of nitrifiers in the oxic water column, no 16S rRNA gene of anammox bacteria was detected in the anoxic waters of the Scharendijke basin. However, these bacteria have been found in the upper two cm of sediment in the Den Osse basin of the same lake, where low sulfide levels were reported (Lipsewers et al., 2016). The absence of anammox in the water column is likely attributed to the highly dynamic conditions in Lake Grevelingen (Egger et al., 2016; van Helmond et al., 2025) and rapid cycles of stratification and mixing. These factors most likely hinder the establishment of anammox bacteria despite the availability of NH_4_^+^ and NO_2_^-^ in the suboxic zone. This may explain why anammox bacteria are more commonly found in systems that are permanently stratified and oxygen-deficient, such as oxygen minimum zones (Kalvelage et al., 2015; Kuypers et al., 2005), the Black Sea (Jensen, et al., 2008), and in anoxic sediments (Lipsewers et al., 2016).

### Factors driving the nitrifying community composition

Both AOA and AOB are found in the water column of the Scharendijke basin, with AOB appearing to be more abundant. This contrasts with the findings for most other marine environments, where AOA usually outcompete AOB (Bouskill et al., 2012; Hutchins & Capone, 2022; Lam et al., 2007; Newell et al., 2013). Even in the North Sea, which exchanges water with marine Lake Grevelingen, archaeal *amoA* gene copy numbers were one or two orders of magnitude higher than those of bacterial *amoA* copies (Wuchter et al., 2006). Furthermore, *Nitrosomonas* species have been successfully enriched from the North Sea water column at an NH_4_^+^ concentration of 500 µM (Haaijer et al., 2013), indicating that AOB may thrive under certain conditions.

In Lake Grevelingen, we observed high concentrations of NH_4_^+^, which might favor AOB over AOA. It is known that AOA have a better affinity for ammonia, with K_s_ values 100-fold lower compared to AOB, so they are often found in oligotrophic systems like the open ocean (Francis et al., 2005; Hatzenpichler, 2012; Jung et al., 2021; Karner et al., 2001; Martens-Habbena et al., 2009; Sakoula et al., 2021). Furthermore, it has been shown that for some AOA, ammonia oxidation rates decrease with increasing NH_4_^+^ concentrations, while AOB ammonia oxidation rates increase (French et al., 2012). Our system is very eutrophic and might therefore preferentially select for AOB instead of AOA in the water column.

Although NH_4_^+^ was detected in oxic waters as early as April, significant removal of NH_4_^+^ was not observed until late August. This suggests that AOB may require not only ammonia but also favorable oxygen conditions to become metabolically active. In the eutrophic Scharendijke basin, competition for oxygen among microbial communities could critically influence the timing and extent of AOB activity. Heterotrophic bacteria, which have high oxygen affinities (Canfield & Kraft, 2022; Kalvelage et al., 2015), may effectively outcompete nitrifiers for oxygen, potentially delaying the onset of nitrification (Cho et al., 2022).

Importantly, Lake Grevelingen is a marine lake with a salinity of 29 to 33 (Żygadłowska et al., 2024) As noted above, AOB are generally more abundant in freshwater systems, while AOA dominate in the marine environment (Bollmann & Laanbroek, 2002; Bouskill et al., 2012; Hutchins & Capone, 2022). This was also shown in a study from the Chesapeake Bay, where in the more inland and freshwater parts, AOB (*Nitrosomonas* and *Nitrosospira*) were dominant, while in the more marine areas, AOA related to *Nitrosopumilus maritimus* took over (Bouskill et al., 2012). Additionally, in the Schelde Estuary, *Nitrosomonas* species were observed but shifted species within the genus when salinity increased (Bollmann & Laanbroek, 2002; De Bie et al., 2001). Thus, AOB can occur in marine systems but are generally not as dominant as AOA (Lam et al., 2007). Still, a few studies show that AOB can outnumber AOA in coastal marine sediments (Bernhard et al., 2007; Freitag et al., 2006; Santoro et al., 2008). Interestingly, it was observed that when AOB are found in the marine water column, these are often *Nitrosospira* species (Bernhard et al., 2007; Freitag et al., 2006; Santoro et al., 2008). However, in our study, we observed that *Nitrosomonas* was the most abundant ammonia oxidizer present and had apparently adapted to the marine salinity of Lake Grevelingen.

Network analysis was used to explore whether the co-occurrence of microbial taxa might indicate interactions or niche partitioning that facilitate the occurrence of AOB. *Nitrosomonadaceae* were strongly correlated with the OM190 and SAR202 lineages, which may reflect shared environmental preferences or indirect interactions. OM190, a lineage within the *Planctomycetota*, has previously been observed in oxic-anoxic transition zones and is present in environments with elevated nitrate concentrations (Storesund et al., 2020; Vitorino et al., 2025). In a meromictic lake in Norway, OM190-affiliated species co-occurred with the archaeal ammonia oxidizer *Nitrosopumilus*, both peaking in abundance at the oxic-anoxic interface (Storesund et al., 2020). Although direct metabolic interaction between the OM190 lineage and *Nitrosomonadaceae* is unlikely, their co-occurrence may result from overlapping niche requirements. For instance, metagenome-assembled genomes (MAGs) affiliated with the SAR202 lineage of the *Chloroflexi* phylum, which, like OM190, display a positive correlation to the *Nitrosomonadaceae* in our analysis, encode a NapAB-type nitrate reductase (Thrash et al., 2017). This indicates a role in nitrate reduction and, therefore, possible usage of nitrate formed by nitrification. However, MAGs affiliated with *Candidatus* Peregrinibacteria, the third lineage that had a positive correlation with the *Nitrosomonadaceae*, did not contain any nitrogen cycling genes, despite occurring in the hypoxic water column of the Gulf of Mexico (Thrash et al., 2017). Therefore, these lineages might have metabolic links beyond nitrogen exchange, such as an exchange of B vitamins, as has recently been demonstrated for *Nitrospina gracilis* in co-culture with heterotrophs (Bayer et al., 2024). Alternatively, these correlations may reflect congruent habitat preferences for ammonium and oxygen. Interestingly, Venetz et al. (2024) reported that methane-oxidizing bacteria remained abundant in the water column of the Scharendijke basin at the end of August but showed signs of reduced methane consumption, indicating a possible inhibition of their activity due to a shift in the microbial community (Venetz et al., 2024).

The main NOB in Lake Grevelingen was *Nitrospina,* which is commonly found in marine and coastal ecosystems (Bristow et al., 2015; Daims et al., 2016; Damashek & Francis, 2018; Mincer et al., 2007), including oxygen minimum zones (Füssel et al., 2012; Sun et al., 2019). We furthermore observed that *Nitrospira* was present in low abundance in March. However, as the presence of *Nitrospira* was not noted in other months, it is probably of minor relevance for nitrite oxidation in the Scharendijke basin, at least during the summer months. In accordance with the fact that *Nitrospina* are more commonly found in areas with low or even without oxygen (Füssel et al., 2012; Sun et al., 2019), the appearance of the NO_2_^-^ and NO_3_^-^ peaks at the oxic-anoxic interface towards the end of August suggests that NO_2_^-^ is oxidized with low oxygen also in the water column of Lake Grevelingen.

## Conclusions

Our study demonstrates that the Scharendijke basin in Marine Lake Grevelingen experienced seasonal ammonium accumulation in 2021 due to restricted vertical mixing caused by strong stratification, with potential additional contributions from lateral ammonium input. Nitrification is only observed in late August and September, when ammonium reaches the oxic-anoxic interface, and is primarily mediated by ammonia- and nitrite-oxidizing members of the *Nitrosomonadaceae* and *Nitrospinaceae* families, respectively, in contrast to the general dominance of AOA in other marine environments. In winter, nitrifiers are largely restricted to the sediment and probably release nitrate into the water column. Despite ammonium being present in the water column from April onwards, oxygen limitation, microbial competition, or other environmental factors may constrain nitrification. These findings emphasize seasonal variations in nitrogen cycling in stratified marine systems, highlighting the need to consider both physical and biological factors in predicting nitrogen conversion rates.

## Acknowledgement

We thank the captain and crew of the R/V Navicula, other NIOZ staff, and all Marix participants for their support during the sampling campaigns. This research was financially supported by a European Research Council Synergy grant MARIX (854088). WL acknowledges support by NWO Veni grant VI.Veni.222.332.

## Conflict of interest statement

The authors declare no competing interests.

## Notes

### Competing Interest Statement

The authors have declared no competing interest.

