## Supplementary Information for "Seasonal dynamics of nitrification in the water column of marine Lake Grevelingen"

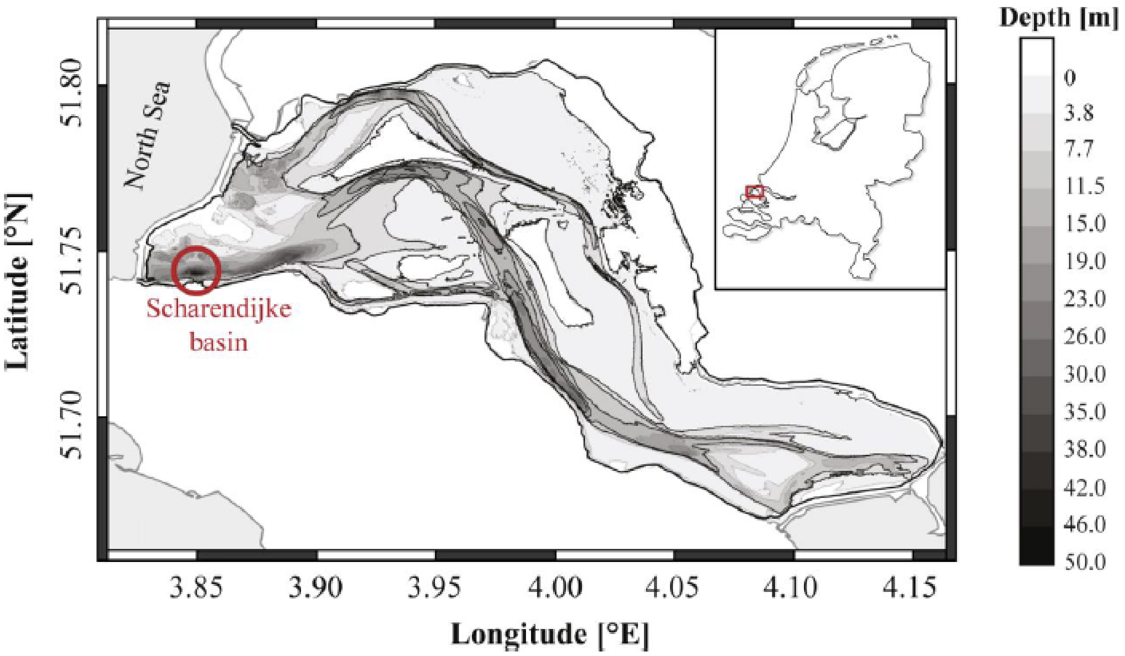


***Figure S1***. *Bathymetry of Lake Grevelingen. The location of the Scharendijke basin (51.742 N; 3.849 E) is indicated by the circle (adapted from Egger et al., 2016).*

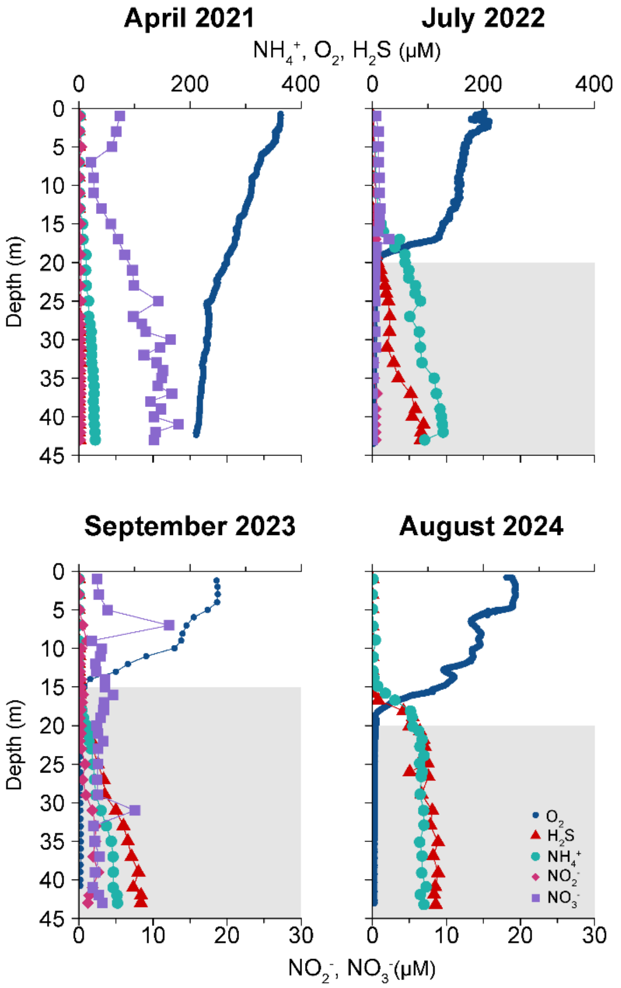


***Figure S2**. Water column profiles of the Scharendijke basin in Lake Grevelingen in 2021 spring, summer 2022, 2023 and 2024. The depth profiles of $O_2$ (blue circle), $H_2S$ (red triangle), $NH_4^+$ (green circle), $NO_2^-$ (red diamond) and $NO_3^-$ (purple square) are depicted. The Grey shaded areas indicate the part of the water column where $O_2$ was below the detection limit of the sensor (< 4 µM).*

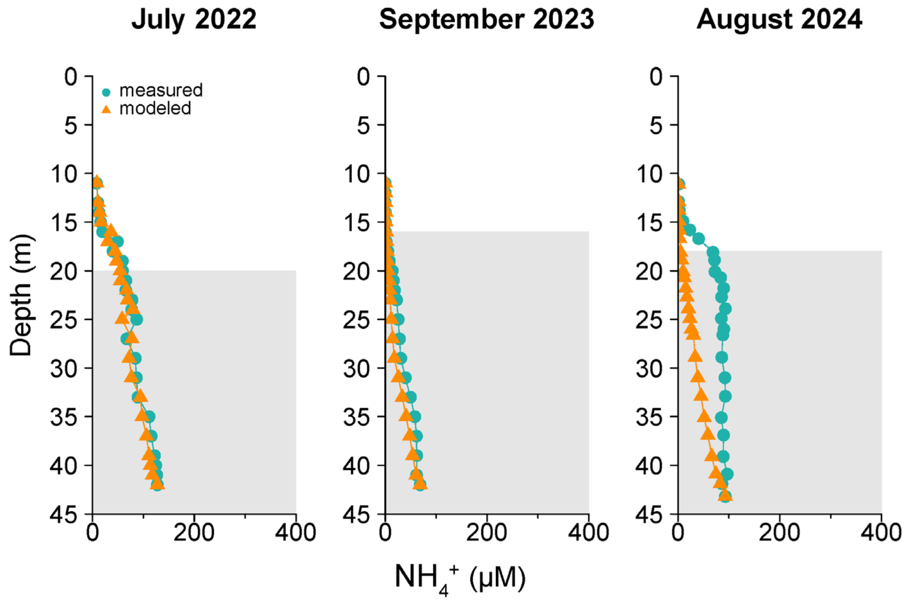


***Figure S3.*** *Water column profiles comparing the measured $NH_4^+$ concentrations with the concentrations predicted by the turbulent diffusion model, calculated for the stratified part of the water column. The samples were obtained in July 2022, September 2023, and August 2024. Measured $NH_4^+$ concentrations are indicated by green circles, the orange triangles indicate the predicted $NH_4^+$ concentrations.*

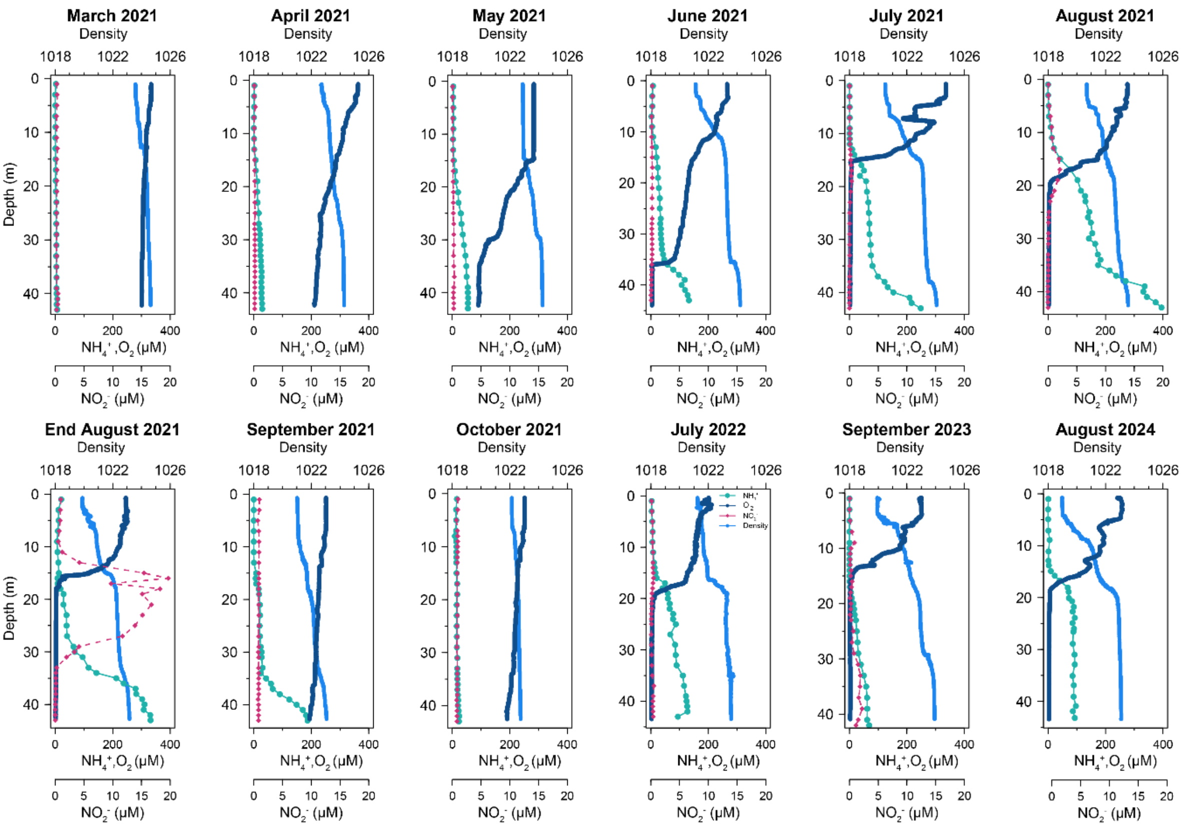


***Figure S4.*** *Depth profiles of the density of the water (in kg/m$^3$, light blue circles), $O_2$ (in µM, dark blue circles), $NH_4^+$ (in µM, light green circles), and $NO_2^-$ (in µM, pink diamonds) of all sampling dates from March till October 2021 and July 2022, September 2023, and August 2024.*

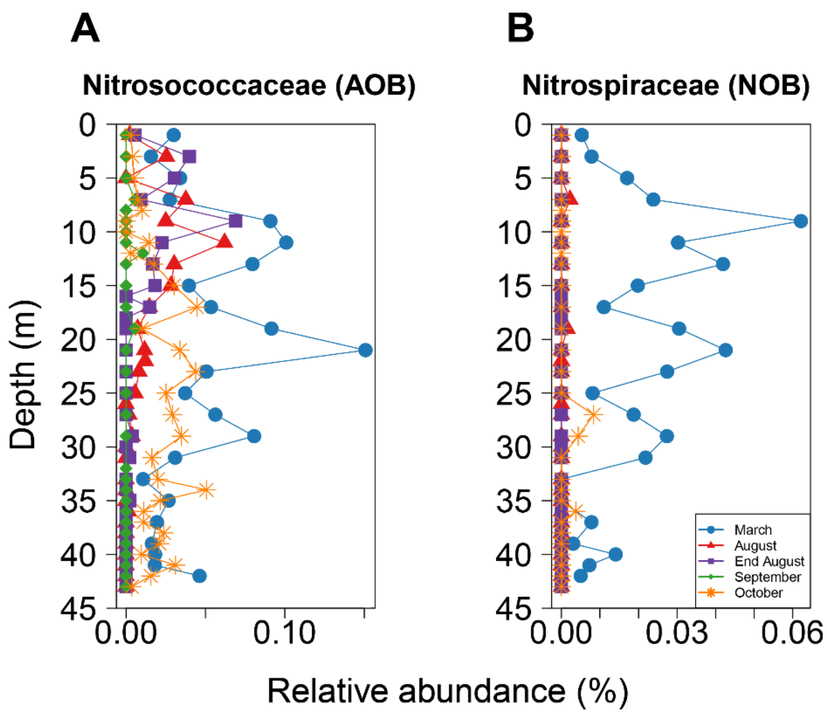


***Figure S5.*** *16S rRNA gene amplicon sequencing-based nitrifier abundance in the Scharendijke basin water column in Lake Grevelingen. Relative abundances of (A) ammonia-oxidizing members of the family Nitrosococcaceae and (B) nitrite-oxidizing Nitrospiraceae in March (blue), August (red), end of August (purple), September (green), and October 2021 (orange).*

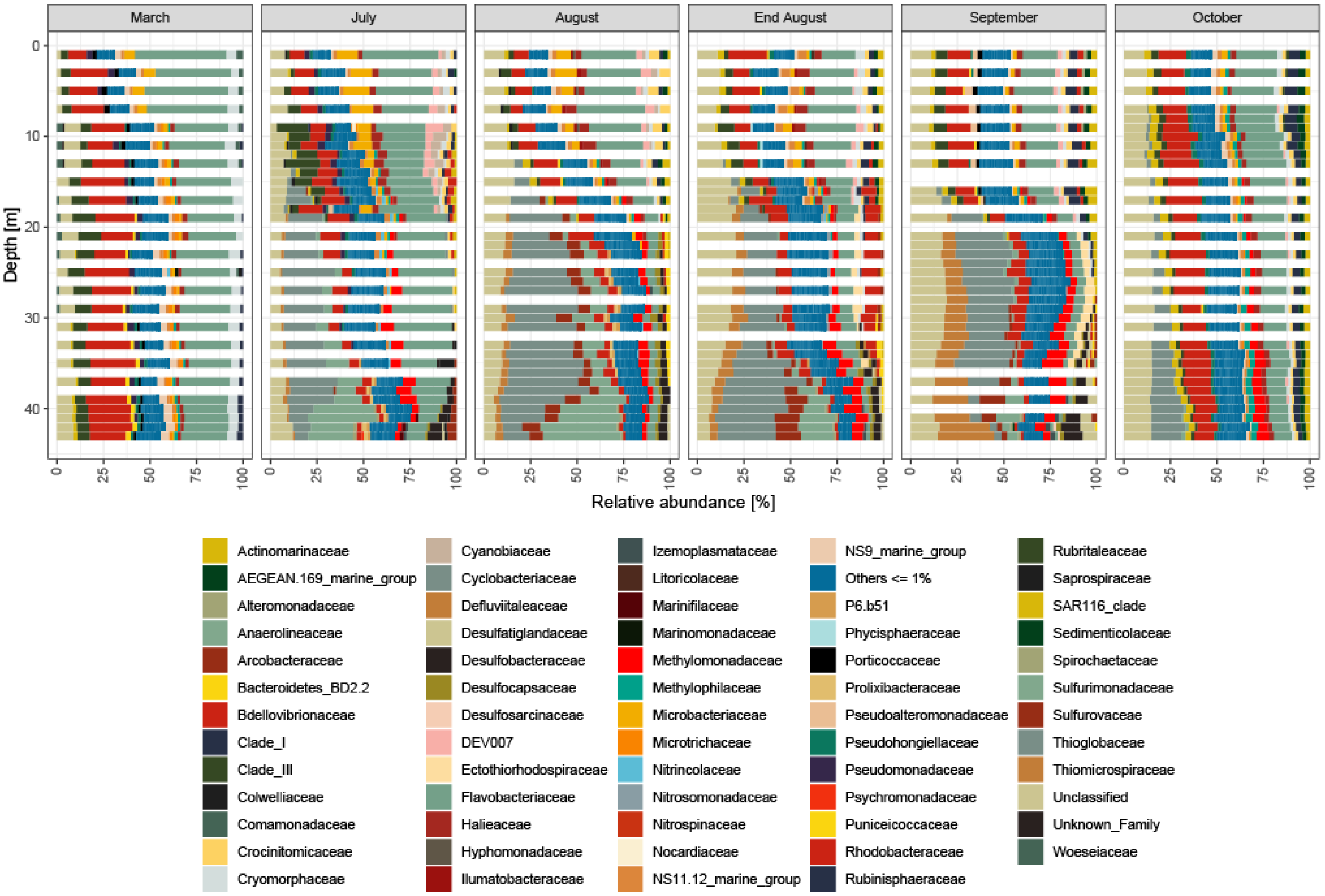


***Figure S6.*** *16S rRNA gene amplicon sequencing-based bacterial relative abundances. Only taxa representing ≥1% of the total bacterial community are shown. The data represent the composition of the bacterial fraction from March to October 2021. Low-abundance taxa (<1%) are grouped as "Others".*

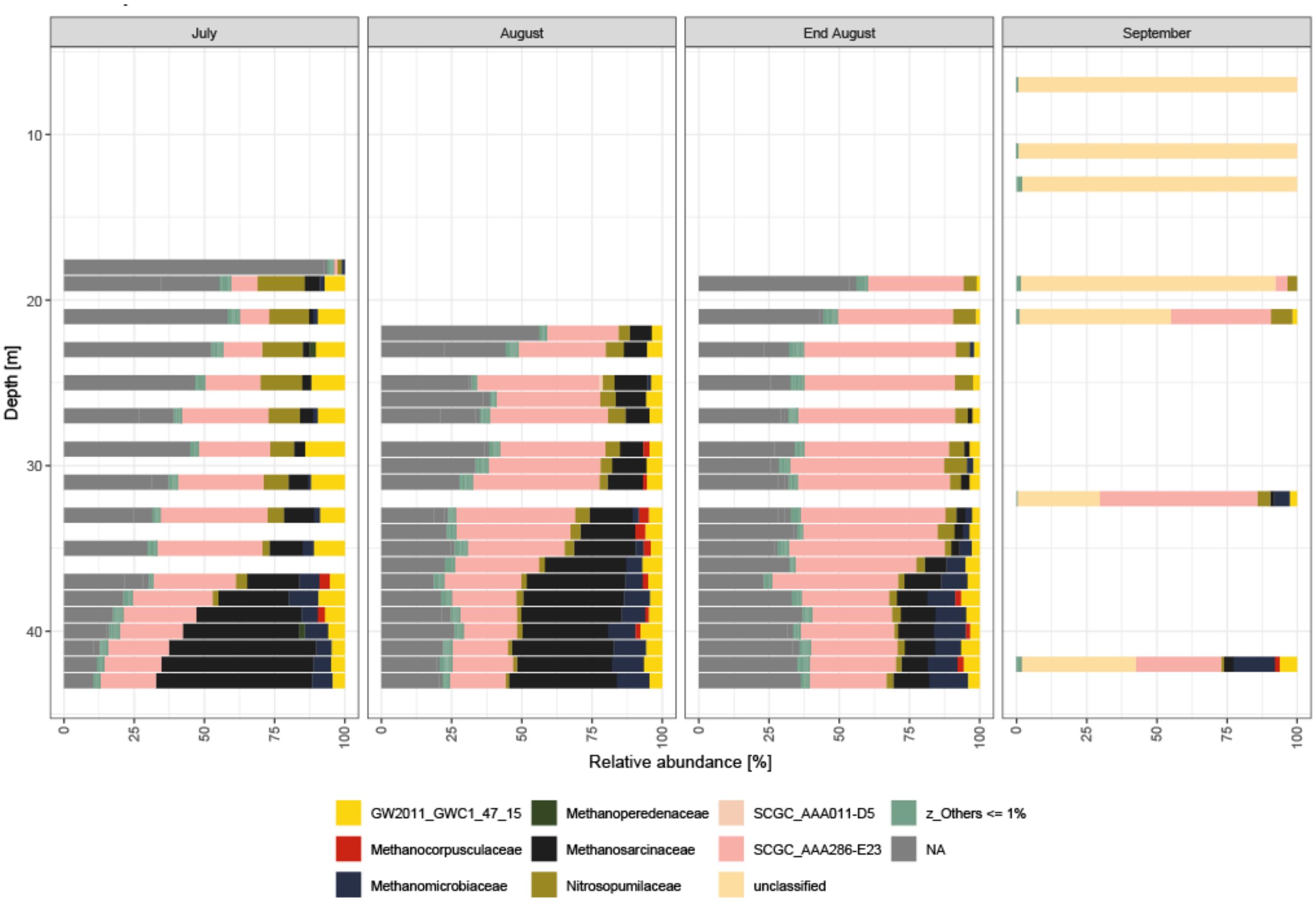


***Figure S7.*** *16S rRNA gene amplicon sequencing-based archaeal relative abundances. Only taxa representing ≥1% of the total archaeal community are shown. The data represent the composition of the archaeal fraction from July to September 2021. Low-abundance taxa (<1%) are grouped as "Others".*

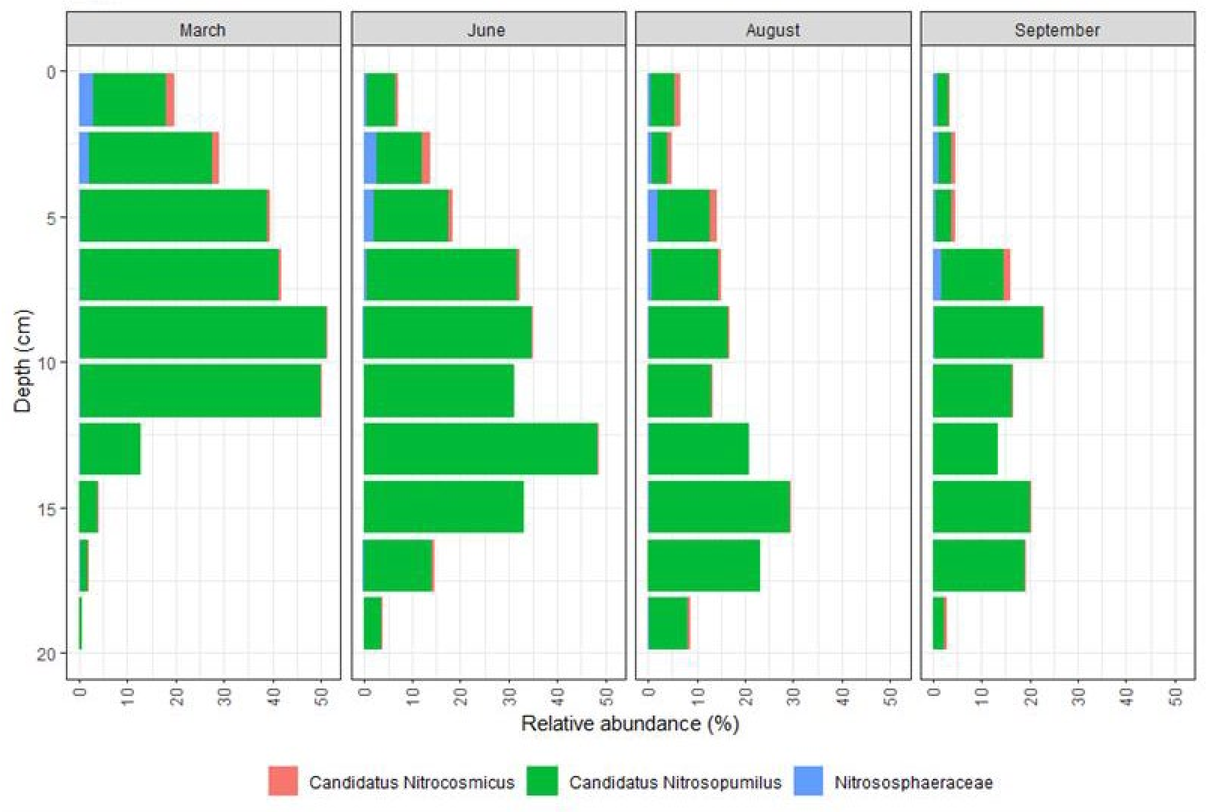


***Figure S8.*** *Relative abundances of archaeal ammonia oxidizers in sediment from the Scharendijke basin in March, June, August, and September 2021. Shown are the relative cumulative relative abundances of members of the Nitrosopumilaceae (green).*

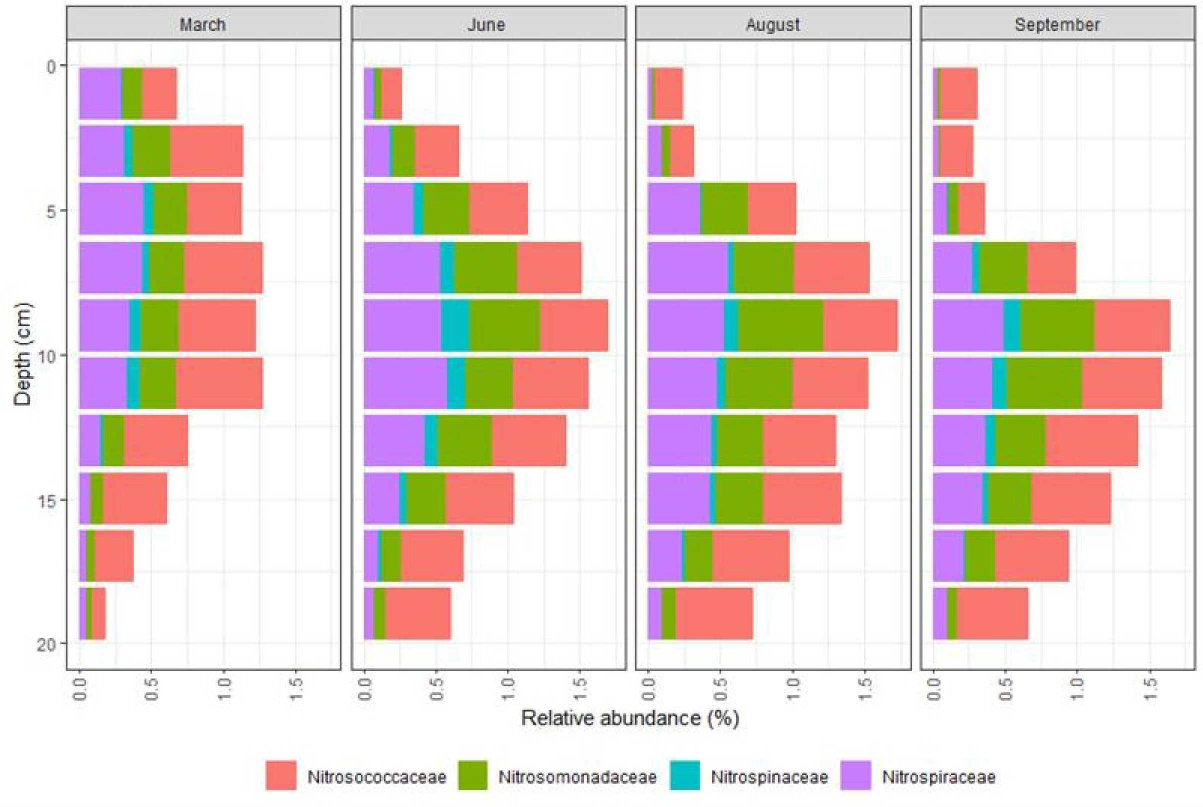


***Figure S9.*** *Relative abundances of bacterial nitrifiers in sediment of Scharendijke basin in March, June, August, and September 2021. Shown are the relative cumulative relative abundances of members of the Nitrosococcaceae (red), Nitrosomonadaceae (green), Nitrospinaceae (turquoise), and Nitrospiraceae (purple).*

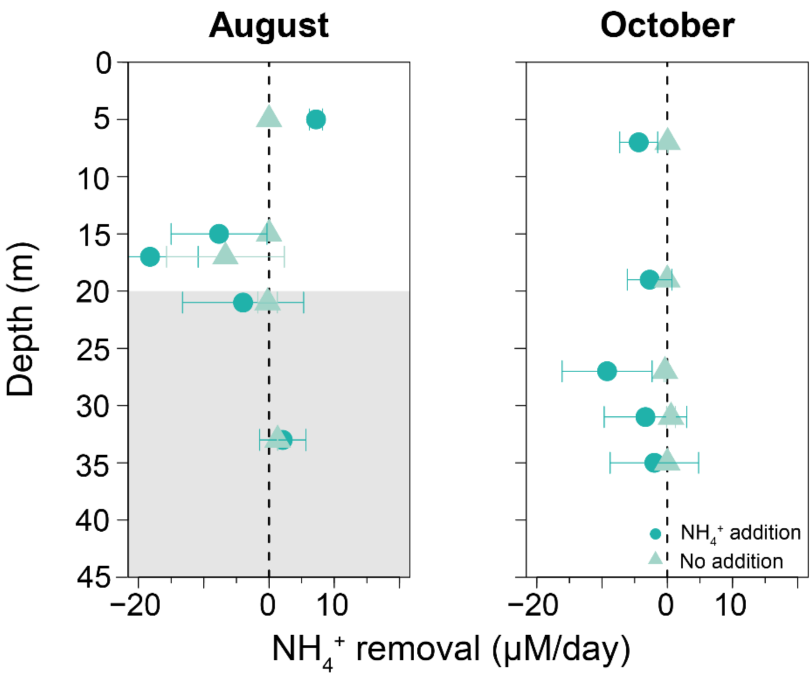


***Figure S10***: *Ammonium removal rates at five selected water depths in the oxic layer, at the start of the oxycline, in the middle of the oxycline, at the oxic-anoxic interface, and in the anoxic layer. Water samples were taken in August and October 2021. $NH_4^+$ removal rates (µM/day) are shown for incubations with (dark blue) and without $NH_4^+$ addition (only endogenous ammonium; light blue). Data points represent the mean ammonium removal rates; the error bars represent the SD based on duplicate measurements. The dashed black line indicates the threshold between net $NH_4^+$ removal (positive values) and potential net production of ammonium via mineralization (negative values). The grey shaded areas indicate the part of the water column where $O_2$ was below the detection limit of the sensor (< 4 µM).*

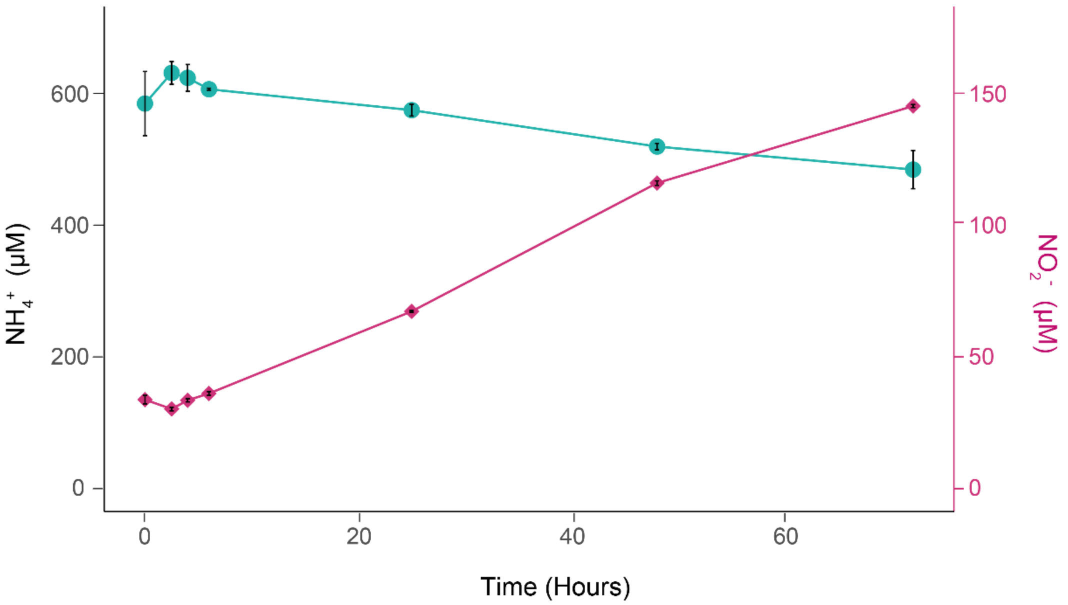


***Figure S11.*** *Representative ammonia oxidation assay showing the conversion of ammonium to nitrite over time. The water sample was taken in August 2021 at the oxic-anoxic interface at a depth of 21 meters in the Scharendijke basin at Lake Grevelingen.* $NH_4^+$ *concentrations (blue/green) are shown on the left, and* $NO_2^-$ *concentrations (pink) on the right y-axis. Data points indicate individual measurements, error bars show the SD based on duplicate measurements.*

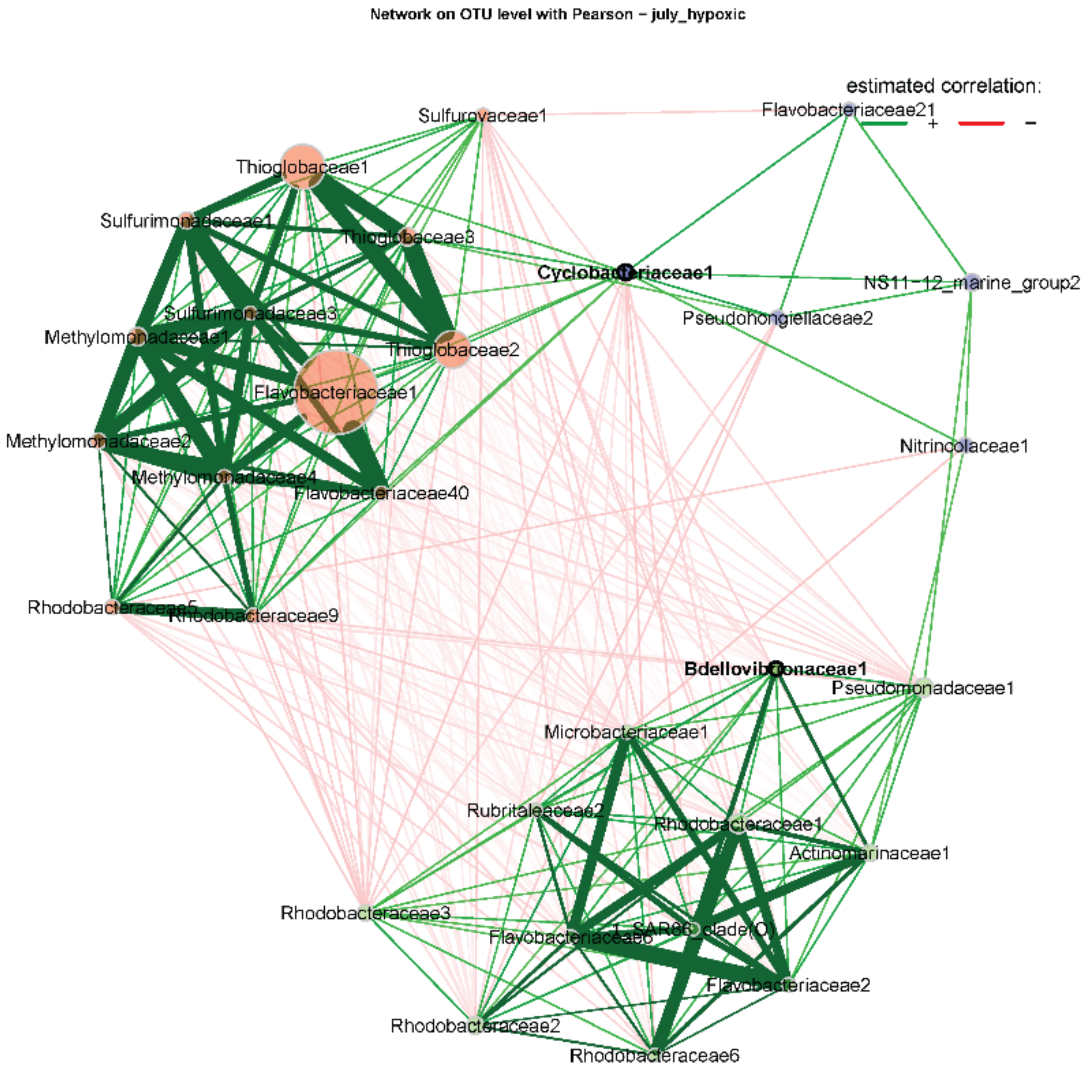


***Figure S12.*** *Co-occurrence networks based on Pearson correlation for the hypoxic (<60 µM $O_2$) area in the water column for July 2021. Edges represent significant correlations, with green lines indicating positive and red lines indicating negative correlations. Stronger positive correlations are emphasized by line width. Node colors indicate distinct microbial clusters, while node size represents eigenvector centrality, highlighting the most influential taxa. Hubs, identified based on betweenness centrality, are highlighted with black node borders and bold font, representing key microbial taxa that serve as connectors between different clusters.*

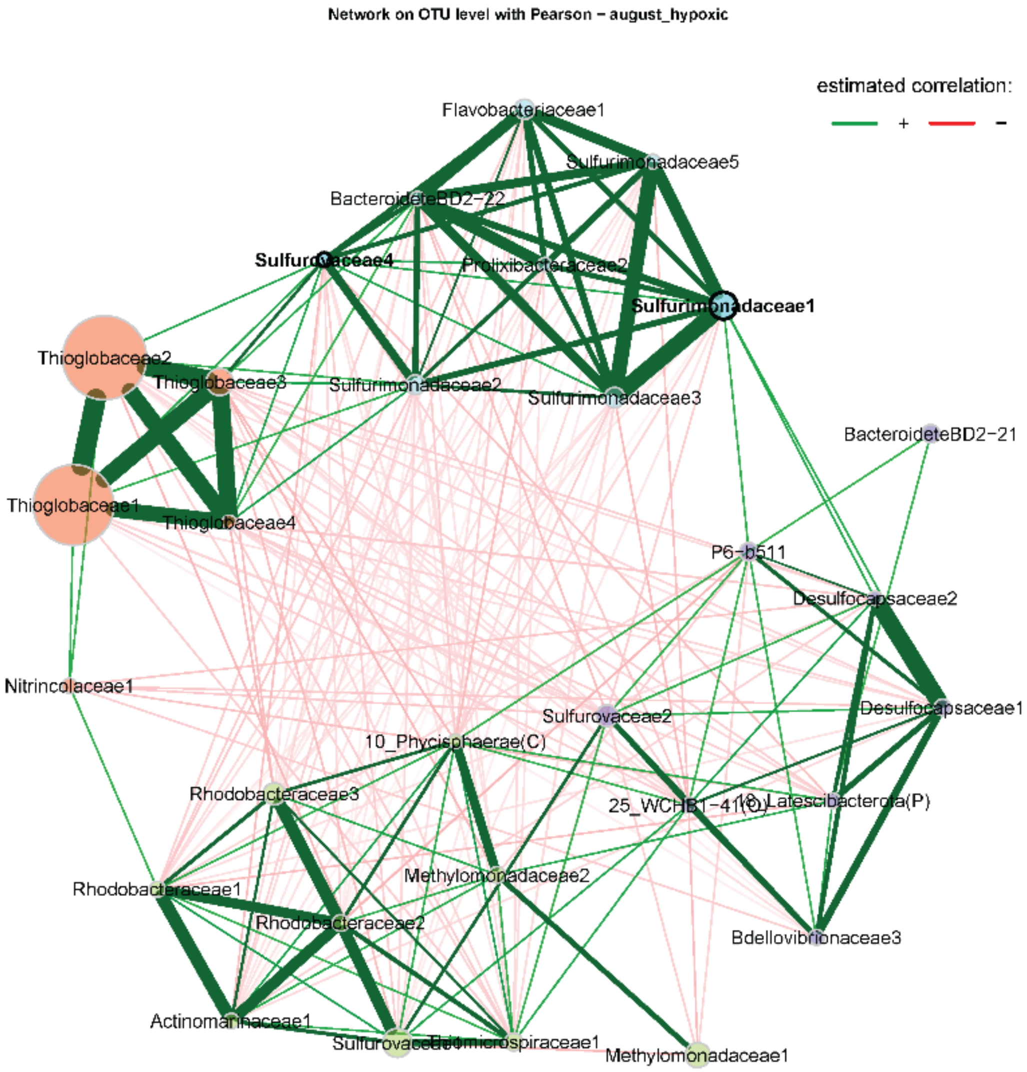


***Figure S13***. *Co-occurrence networks based on Pearson correlation for the hypoxic (60 µM $O_2$) area in the water column for August 2021. Edges represent significant correlations, with green lines indicating positive and red lines indicating negative correlations. Stronger positive correlations are emphasized by line width. Node colors indicate distinct microbial clusters, while node size represents eigenvector centrality, highlighting the most influential taxa. Hubs, identified based on betweenness centrality, are highlighted with black node borders and bold font, representing key microbial taxa that serve as connectors between different clusters.*

***Table S1.*** *Water column sampling dates in 2021, 2022, 2023, and 2024*

| Year | Sample name | Sampling day |
|---|---|---|
| 2021 | March 2021 | March17 |
| | April 2021 | April 21 |
| | May 2021 | May 5 |
| | June 2021 | June 5 |
| | July 2021 | July 12 |
| | August 2021 | August 11 |
| | End August 2021 | August 31 |
| | September 2021 | September 20 |
| | October 2021 | October 12 |
| 2022 | July 2022 | July 31 |
| 2023 | September 2023 | September 13 |
| 2024 | August 2024 | August 14 |

***Table S2****. Integrated water column* $NH_4^+$ *concentrations (in mmol* $m^{-2}$*) in the middle (10-35 m) and bottom layers (35-45 m) of the Scharendijke basin calculated for the samples from June, July, August, End August, and September 2021 adapted from Zygadlowvska et al., 2024.*

| Water column layer | June | July | August | End August | September |
|---|---|---|---|---|---|
| Middle | 716 | 903 | 2819 | 1271 | 355 |
| Bottom | 950 | 1459 | 2873 | 2762 | 1570 |

***Table S3****. Metagenomic read-based relative abundances of archaea and bacteria at 4 depths in the water column of September 2020.*

| Depth [m] | Archaea [%] | Bacteria [%] |
|---|---|---|
| 25 | 4 | 96 |
| 32 | 1 | 99 |
| 35 | 4 | 96 |
| 42 | 1 | 99 |

***Table S4.*** *Metagenomic read-based relative abundances of archaea and bacteria at 3 depths in the sediment of September 2020.*

| Depth [cm] | Archaea [%] | Bacteria [%] |
|---|---|---|
| 0-2 | 2 | 98 |
| 9-11 | 2 | 98 |
| 15-17 | 3 | 97 |
